# cAMP signalling is required for the actions of IP_3_ on Ca^2+^-transients in cardiac atria and beating rate in sino-atrial node

**DOI:** 10.1101/694349

**Authors:** Rebecca A Capel, Thomas P Collins, Samuel J Bose, Skanda Rajasundaram, Thamali Ayagama, Manuela Zaccolo, Rebecca AB Burton, Derek A Terrar

**Affiliations:** Department of Pharmacology, British Heart Foundation Centre of Research Excellence, University of Oxford, Mansfield Road, Oxford, Oxon, OX1 3QT; Department of Physiology, Anatomy and Genetics, University of Oxford, Parks Road, OX1 3PT

## Abstract

Inositol trisphosphate (IP_3_) is a major Ca^2+^-mobilising second messenger and atrial IP_3_ receptor (IP_3_R) expression is greatly increased in atrial fibrillation (AF). Cardiac atrial and sino-atrial node (SAN) myocytes also express Ca^2+^-stimulated adenylyl cyclases (AC1 and AC8); however the pathways underlying atrial AC1 and AC8 activation are unknown. We investigated whether IP_3_ signalling in cardiac atria and SAN utilises ACs. Immunocytochemistry in isolated guinea pig atrial myocytes identified co-localisation of type 2 IP_3_Rs with AC8, while AC1 was located in close vicinity. UV photorelease of IP_3_ significantly enhanced Ca^2+^ transient amplitudes following stimulation of atrial myocytes (31 ± 6 % increase 60 s post photorelease, n=16), an effect abolished by inhibitors of ACs (MDL-12,330) or PKA (H89). The maximum rate change observed in spontaneously-beating murine right atrial preparations exposed to phenylephrine (14.7 ± 0.5 %, n=10) was significantly reduced by 2.5 μmol/L 2-APB and abolished by a low dose of MDL-12,330. These observations are consistent with a functional interaction between IP_3_ and cAMP signalling involving Ca^2+^ stimulation of ACs in cardiac atria and the SAN. Structural evidence supports AC8 as the most likely effector. This signal transduction mechanism is important for future study in atrial physiology and pathophysiology, particularly AF.

## Introduction

Calcium handling in the heart is vital to normal physiological function, arising from the interaction of multiple, highly regulated, calcium signalling pathways. The atrial and ventricular chambers of the heart have very different functions and therefore it is not surprising that there are many differences between atrial and ventricular myocytes in excitation-contraction coupling and in the handling of Ca^2+^ ions by different intracellular compartments. One characteristic feature of atrial myocytes is the relative abundance of receptors for inositol trisphosphate (IP_3_) compared with ventricular myocytes ^1^. IP_3_ is a Ca^2+^-mobilising second messenger ^2^ which acts to open IP_3_ receptors (IP_3_R), located on the sarcoplasmic reticulum (SR) of cardiomyocytes ^1,3^. IP_3_ is positively inotropic in atrial ^1^ and ventricular ^4^ preparations, and is positively chronotropic in the sino-atrial node ^5,6^. IP_3_ is synthesised upon stimulation of phospholipase C (PLC), commonly but not exclusively by G-protein coupled receptors associated with Gq ^7^. In cardiac myocytes endothelin-1 (ET-1), angiotensin II (Ang-II) and phenylephrine (PE) all increase intracellular IP_3_ level ^8^ via their actions at the Gq-coupled ET-A, Ang-II and α-adrenergic receptors respectively.

Early functional studies revealed a much greater effect of IP_3_-associated stimuli on the contractility of atrial preparations than upon their ventricular counterparts ^9^ and expression of IP_3_R type 2 (IP_3_R2) is now known to be at least six times greater in atrial myocytes ^1^. IP_3_R expression is significantly increased during atrial fibrillation in both human patients ^10^ and animal models ^11^. Further, inhibiting G_q_-coupled AngII receptors has been shown to prevent the early remodelling associated with rapid atrial pacing ^12^. Patently, this evidence demonstrates that understanding the functions and underlying physiology of the IP_3_ pathway is particularly important in the cardiac atria.

In healthy atrial myocytes, Gq-associated signalling causes an IP_3_R-dependent increase in the Ca^2+^ spark rate of quiescent myocytes and amplitude of the stimulated Ca^2+^ transient and Ca^2+^ current ^13^, effects matched on direct application of IP_3_ ^1,14^. Interestingly, even in healthy cells, IP_3_-dependent stimulation can be associated with the generation of spontaneous diastolic Ca^2+^ events ^1,3^.

Adenylyl cyclase (AC) enzymes catalyse the production of cAMP. cAMP, in turn, activates PKA and, in the sino-atrial node, also directly regulates the funny current I(f) ^15^. AC5/6 are the predominant AC isoforms traditionally associated with cardiac myocytes ^16^, but atrial and sino-atrial node myocytes also express the Ca^2+^-stimulated isoforms AC1 and AC8 ^17^. Chelation of intracellular Ca^2+^ using BAPTA, or inhibition of ACs using MDL-12,330 (MDL), reduces I(f) in guinea pig sino-atrial node myocytes by shifting the voltage of half-activation to more hyperpolarised voltages; an effect consistent with changes in cellular cAMP. This effect of BAPTA on I(f) is reversed by direct stimulation of ACs using forskolin but is not potentiated by further inhibition of ACs using MDL, consistent with the hypothesis that cAMP from Ca^2+^-dependent ACs affects I(f) under physiological conditions in cardiac pacemaker cells ^17^. Similarly, in guinea pig atrial myocytes BAPTA and MDL reduce peak I_CaL_ amplitude and the effect of BAPTA is abolished when it is applied in the presence of forskolin or high concentrations of patch-applied cAMP ^18^. The effects of BAPTA appear to be calmodulin, but not CaMKII, dependent ^19^, consistent with the known Ca^2+^-dependent activation mechanism of ACs 1 and 8 ^20,21^. Further, expression of Ca^2+^-stimulated AC1 enhances beating rate in HCN2-mediated ‘biological pacemakers’ ^22^.

Considering the above lines of evidence, we sought to investigate the importance of cellular cAMP generation to the effect of IP_3_-dependent signals in the atria and sino-atrial node.

## Results

### Type 2 IP_3_ receptors are co-localised with AC8 in cardiac atrial myocytes

In agreement with published literature ^1^, type 2 IP_3_ receptors were visualised in a punctate pattern at the cell periphery consistent with a position on junctional SR (Figure 1B). Staining for type 1 (Figure 1A) and type 3 IP_3_ (Figure 1C) receptors did not demonstrate a distinct sub-cellular pattern.

**Figure 1.**
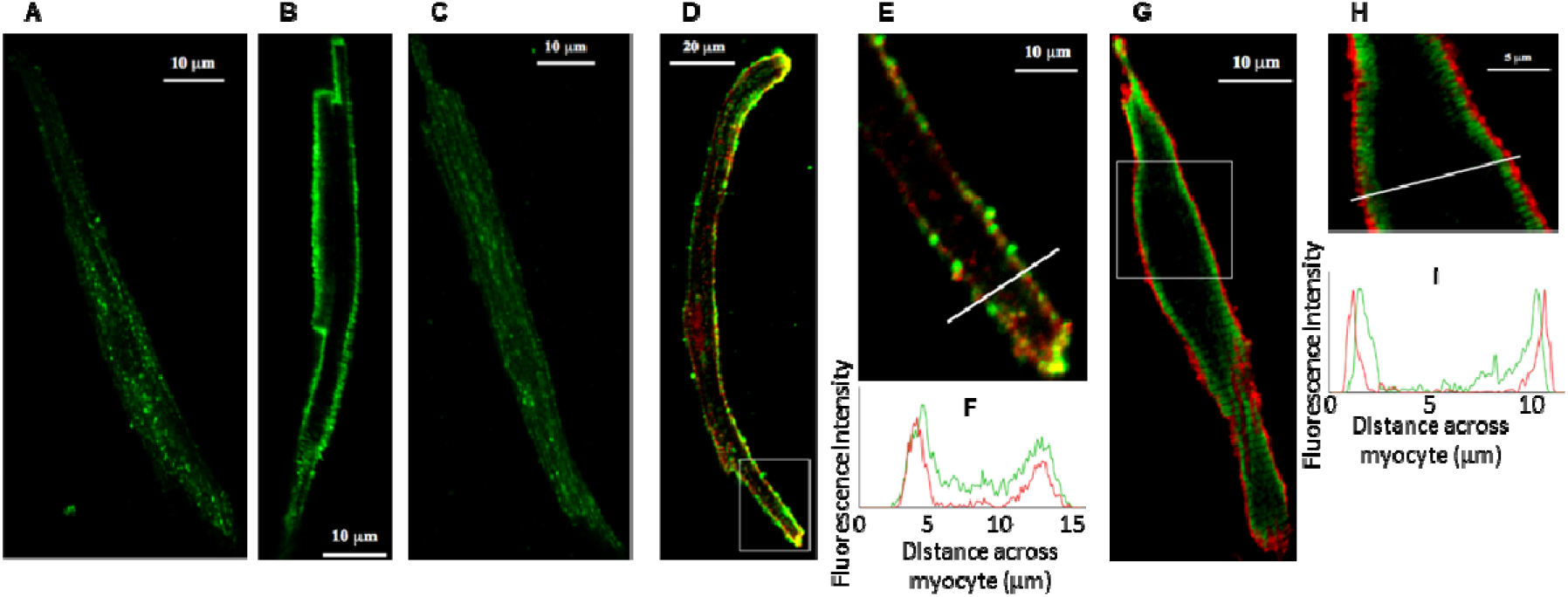
IP_3_ Receptor Type 2 Co-localises with Adenylyl Cyclase 8 in guinea pig atrial myocytes. **A - C.** Representative examples of fixed, isolated guinea pig atrial myocytes labelled for **A.** IP_3_R1, **B.** IP_3_R2 and **C.**IP_3_R3. **D.** Representative example of a fixed, isolated guinea pig atrial myocyte co-immunolabelled for IP_3_R2 (red) and AC8 (green). **E.** Digital zoom of the area indicated on image D. **F.** Intensity plot to show staining intensity along the line shown in E. Full dataset Pearson overlap coefficient R = 0.81 ± 0.02 (n=14). **G.** Representative example of a fixed, isolated guinea pig atrial myocyte co-immunolabelled for IP_3_R2 (red) and AC1 (green). **H.** Digital zoom of the area indicated on image G. **I.** Intensity plot to show staining intensity along the line shown in H. Full dataset Pearson overlap coefficient R = 0.36 ± 0.03 (n=18).

As has been previously described ^17,18^, immunolocalisation of AC8 indicated a band at or just beneath the sarcolemma. Pixel by pixel analysis revealed substantial co-localisation between AC8 and type-2 IP_3_ receptors in isolated guinea pig atrial myocytes, Pearson overlap coefficient R = 0.81 ± 0.02 (n=14, Figure 1D-F).

AC1 staining was localised to a band which was consistently nearby but predominantly just inside type 2 IP_3_ receptors and signals were not substantially overlapping (R = 0.36 ± 0.03, n=18, Figure 1G-I).

### The effect of IP_3_ on cellular Ca2+ transients requires functional adenylyl cyclases and PKA

IP_3_ is not cell permeant and is broken down rapidly within cells. In addition, as activation of α-ARs (e.g. using PE) may result in signalling via alternative pathways including activation of PKC via diacylglycerol (DAG) ^23^, for our experiments we used a cell-permeant, caged version of the compound (IP_3_/PM) to provide cell stimulation specifically via this second messenger from an exogenous source. This IP_3_ compound crosses the cell membrane, is de-esterified by constitutive esterase activity and trapped, and finally can be activated by ‘uncaging’ through brief exposure to UV light.

If Ca^2+^ release through IP_3_ receptors is stimulating cAMP production and PKA activity through AC8, and/or AC1, which is plausible given the immunocytochemistry described in Figure 1, then both functional adenylyl cyclases and PKA would be required for IP_3_ to have a complete effect in cardiac atrial cells. Guinea pig atrial myocytes exhibited the classical ‘U-shaped’ activation pattern of cellular Ca^2+^ transient (Figure 2A). Photorelease of IP_3_ in isolated cardiac atrial myocytes led to a gradual increase in stimulated Ca^2+^ transient amplitude (e.g. 31 ± 6 % increase 60 s post photorelease, n=16, Figure 2B+C). This response was completely abolished in the presence of either the adenylyl cyclase inhibitor MDL (3 μmol/L, n=6, Figure 2B+D), or PKA inhibitor H89 (1 μmol/L, n=9, Figure 2B+E), e.g. change in Ca^2+^ transient at 60 s post photorelease of −9 ± 2 % in the presence of MDL and −16 ± 8 % in the presence of H89. The control IP_3_ response was significantly greater than that of MDL or H89 at all measured timepoints after IP_3_ photorelease (*P*<0.05, ANOVA), whilst the responses seen in the presence of MDL or H89 were not significantly different from one another throughout all timepoints (*P*>0.05, ANOVA).

**Figure 2.**
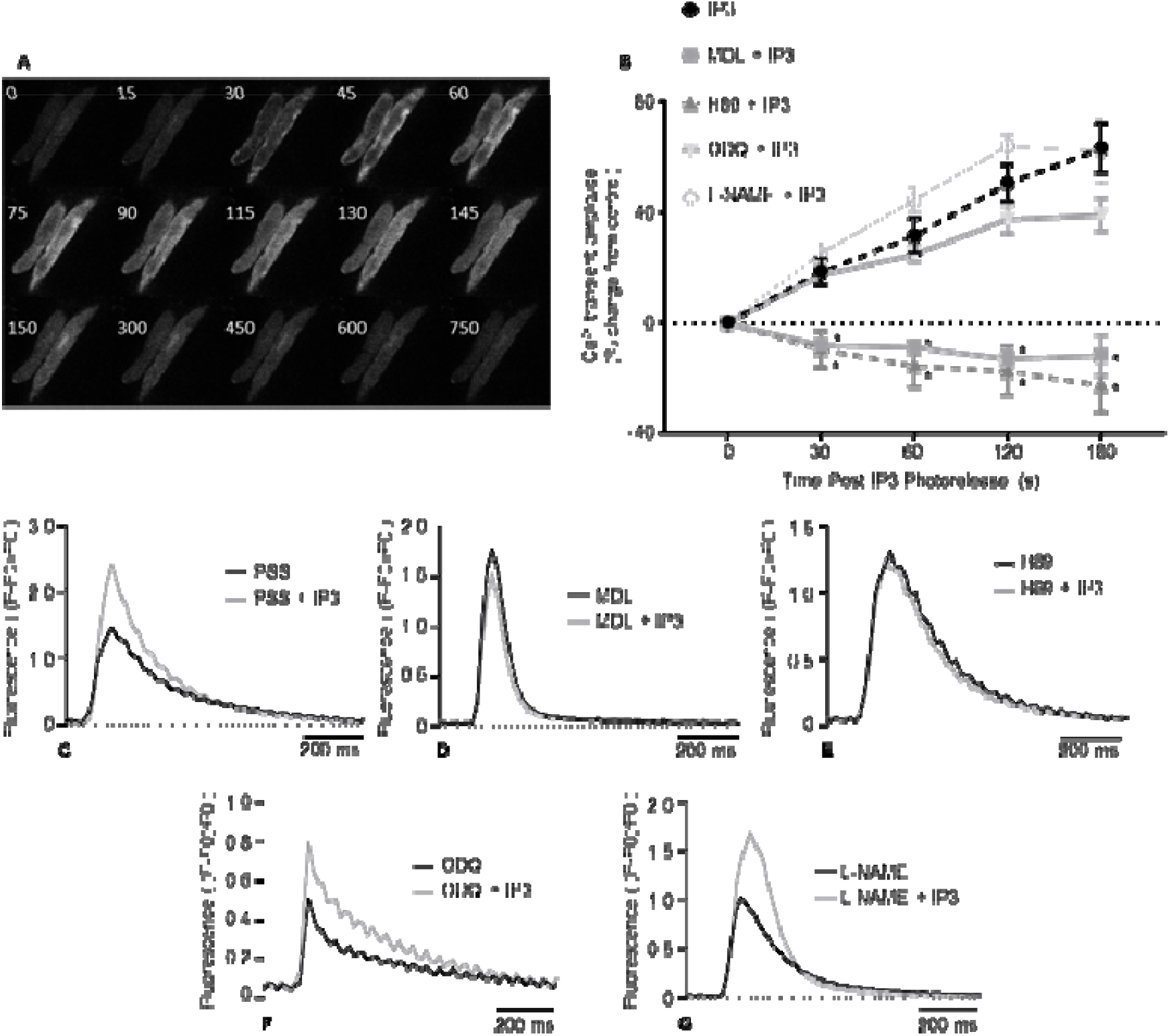
The direct actions of IP_3_ in guinea pig atrial myocytes require adenylyl cyclases and PKA. **A.**Montage to show the progress of a control Ca^2+^ transient in PSS illustrating the classical U-shaped activation pattern. Numbers indicate time in ms from the start of the recording. **B.** Summary data to show cellular responses to IP_3_ photorelease (0.5 μmol/L caged-IP_3_/PM loaded for 1 h) in isolated guinea pig atrial myocytes under control conditions (n=16) and in the presence of 1 μmol/L H89 (n=9), 3 μmol/L MDL (n=6), 100 μmol/L L-NAME (n=4) and 10 μmol/L ODQ (n=10). * denotes significant difference in comparison to IP_3_ photorelease alone (*P* < 0.05, ANOVA with Dunnet’s post-hoc test). **C-G.** Representative Ca^2+^ transient under control conditions and 120 s post photorelease of IP_3_ in: **C.** PSS, **D.** MDL, **E.** H89, **F.** ODQ and **G.** L-NAME.

PE responses in cat atrial myocytes have been reported to be dependent on nitric oxide modulated soluble guanylyl cyclase activity ^24^. We therefore carried out IP_3_ photorelease in the presence of either 10 μmol/L ODQ to inhibit soluble guanylyl cyclase, or 100 μmol/L L-NAME to inhibit nitric oxide synthase. There was no change in the response to IP_3_ photorelease in the presence of ODQ (P>0.05, ANOVA, n=10, Figure 2B+F) or L-NAME (P>0.05, ANOVA, n=4, Figure 2B+G); under both conditions Ca^2+^ transient amplitude increased significantly over time, beginning rapidly after photorelease of IP_3_, and was not significantly different to control at any timepoint.

### The positive chronotropic effect of PE on the sino-atrial node also requires functional adenylyl cyclases

It has been established that endogenous generation, or exogenous administration, of IP_3_ in the sino-atrial node leads to an increase in spontaneous beating rate, accompanied by an increase in Ca^2+^ transient amplitude ^5^, whilst cAMP from Ca^2+^-stimulated adenylyl cyclases has been shown to modulate the funny current in these cells ^17^. Spontaneously beating atrial tissue preparations can also provide a measure of sino-atrial node activity through measurement of beating rate. Log(concentration)-response curves to PE in the concentration range 0.1 – 30 μmol/L were carried out on spontaneously beating isolated murine right atria in the presence of 1 μmol/L metoprolol to ensure no confounding action of β-adrenergic receptors. Under these conditions, the positive chronotropic response to PE fit a standard agonist dose-response curve with an EC50 of 1.12 μmol/L (95 % CI 0.56 to 2.22) and a maximum rate increase of 15.1 ± 0.2 % (n=10, Figure 3A).

**Figure 3.**
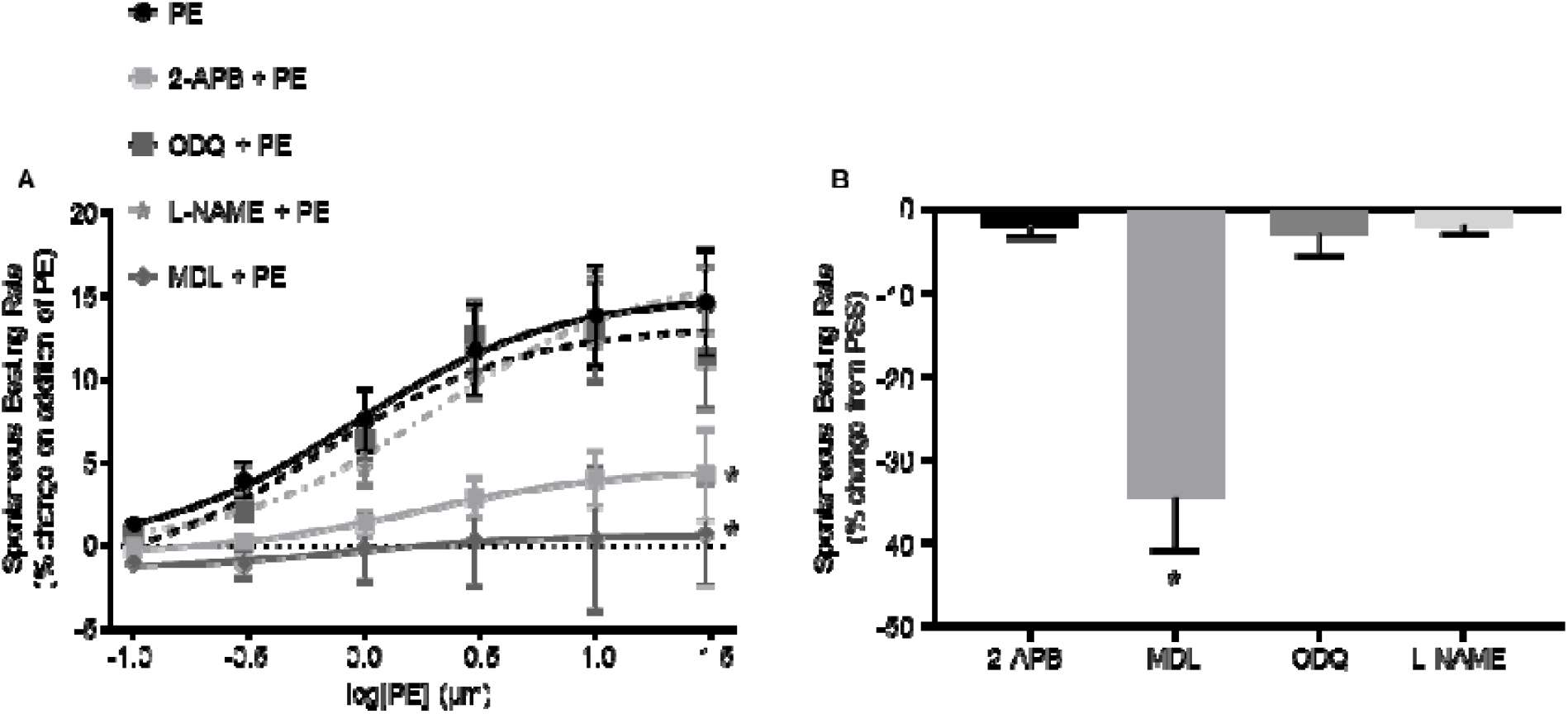
The positive chronotropic effect of PE requires adenylyl cyclases. **A.** Dose-response curves to show the change in beating rate on cumulative addition of PE to spontaneously beating murine right atrial preparations under control conditions (n=10) and in the presence of either 2-APB (2.5 μmol/L, n=7), MDL (1 μmol/L, n=5), L-NAME (100 μmol/L, n=6) or ODQ (30 μmol/L, n=5). * denotes significant reduction in maximum response of the fitted curve by F-test. **B.** Comparison of beating rate change in spontaneously beating murine atrial preparations from control (stable beating in physiological salt solution) on addition of the inhibitors used in A, prior to stimulation by PE. * denotes significant reduction from control by Paired t-test with Bonferroni correction.

Addition of a low concentration of 2-Aminoethyl diphenylborinate (2-APB) (2.5 μmol/L), which is low enough to inhibit IP_3_-dependent effects in cardiomyocytes without altering cellular Ca^2+^ transient amplitude or SERCA function ^5,25–27^, had no significant effect on right atrial beating rate over the course of at least 30 min (P>0.05, Paired t-test, n=7, Figure 3B). In the presence of 2-APB, the maximum rate increase observed on addition of PE was significantly reduced, to 4.7 ± 0.2 % (n=7, Figure 3A), without significant effect on EC50 (1.69 μmol/L, 95 % CI 0.99 to 2.89).

Addition of 1 μmol/L MDL to inhibit adenylyl cyclase activity led to a 34.5 ± 6.4 % reduction in beating rate in the absence of further intervention (Figure 3B, *P*<0.05, Paired t-test, n=5). Under these conditions, bath application of cumulative doses of PE no longer led to an increase in beating rate (maximum rate change 0.7 ± 0.2 %, n=5, Figure 3A). In agreement with the IP_3_ photorelease data detailed above, neither L-NAME (100 μmol/L, n=6), nor ODQ (30 μmol/L, n=5) had a significant effect upon spontaneous beating rate under control conditions (Figure 3B), the maximum response to PE or the EC50 of the response to PE (Figure 3A).

## Discussion

This study represents the first measurements that link direct cellular stimulation with IP_3_ in atrial myocytes to downstream actions via the generation of cAMP and activation of PKA. Our work is consistent with the hypothesis that interaction of IP_3_-mediated Ca^2+^ release with the cAMP system is essential for the positive inotropic and chronotropic effects of this compound in the cardiac atria and sino-atrial node, and that this is physiologically important in the response of these tissues to α-adrenoceptor stimulation. Structural studies using immunostaining methods, which initially led us to investigate this intriguing possibility within our preparations, highlight the Ca^2+^-stimulated isoform AC8 as a probable candidate for this interaction, although involvement of AC1 cannot be excluded.

Ten mammalian adenylyl cyclase isoforms have been discovered, nine membrane bound and one soluble form. Of these, three are Ca^2+^-sensitive; AC1 is CaM-dependently Ca^2+^ stimulated ^20^ with an EC50 for Ca^2+^ of 75 nmol/L ^28^, AC8 is CaM-dependently stimulated ^21,29^ with a Ka for Ca^2+^ activation of ~0.5 μmol/L ^30^ and AC5 is CaM-independently inhibited ^31,32^. The majority of previous studies on AC1 and AC8 pertain to roles in the brain, where these enzymes have been implicated in a range of processes including spatial memory formation ^33–35^, neurodevelopment ^36^, responses to inflammatory pain ^37^ and opioid dependence ^38^. AC1 may also have a role in podocytes of the glomerulus of the kidney ^39^. Our immunocytochemistry data demonstrates that AC8 is found in close proximity to IP_3_Rs in cardiac atrial myocytes whereas AC1 is found in a band just inside these receptors. AC8, therefore, is ideally positioned to transduce local changes in Ca^2+^ into the cAMP-dependent and PKA-dependent effects detailed in this paper; namely the modulation of cellular Ca^2+^ transients in response to IP_3_. Given the known position of IP_3_Rs on the junctional SR ^1,3^, it is not possible for our staining to distinguish whether AC8 is located on the SR itself or on the surface membrane, situated less than 20 nm away ^40,41^. Sucrose-based fractionation of isolated SAN myocytes has indicated that AC1 and AC8 activity is most associated with fractions also containing caveolin-3 ^42^. In other cell types AC8 has been localised to caveolae ^43^, and disruption of lipid rafts has been seen to abolish the stimulation of this cyclase by Ca^2+^ ^30^. Taken together, this evidence is consistent with a surface membrane distribution of this enzyme. Although it seems most likely that Ca^2+^ released via IP_3_Rs activates colocalised AC8, the possibility that this Ca^2+^ could also diffuse to activate nearby AC1 cannot be excluded.

The schematic shown in Figure 4 provides an illustration of the cellular pathway, supported by the data presented in this paper, which may result following activation of the IP_3_ signalling cascade. Increased generation of cAMP via Ca^2+^ activation of AC8 (or AC1) may lead to activation of PKA and multiple downstream actions that result in increases in cellular calcium transient amplitude or beating rate at the sino-atrial node. In particular, previous publications from the Terrar group support a role for AC8/AC1 in modulation of: I(f) in sino-atrial node myocytes ^17^ and L-type Ca^2+^ current in atrial myocytes ^18^. The precise contributions of these actions require further investigation and may also include PKA mediated phosphorylation of ryanodine receptors (RyR) ^44^, L-type Ca^2+^ channels (LTCC) ^45^, phospholamaban (PLB) ^46^ and the Na^+^/Ca^+^ exchanger (NCX) ^47,48^. It is possible that the activity of PKA augments regulation by PKC, which has also been well documented at these same target sites and is similarly activated via stimulation of G_q_ coupled receptors ^49–53^. The use of caged-IP_3_ in this study however rather than stimulation of G_q_ coupled receptors (Figure 2) demonstrates that the effects on cellular Ca^2+^ observed in the present study can occur via the effects of IP_3_ signalling specifically and are independent of activation of DAG.

**Figure 4.**
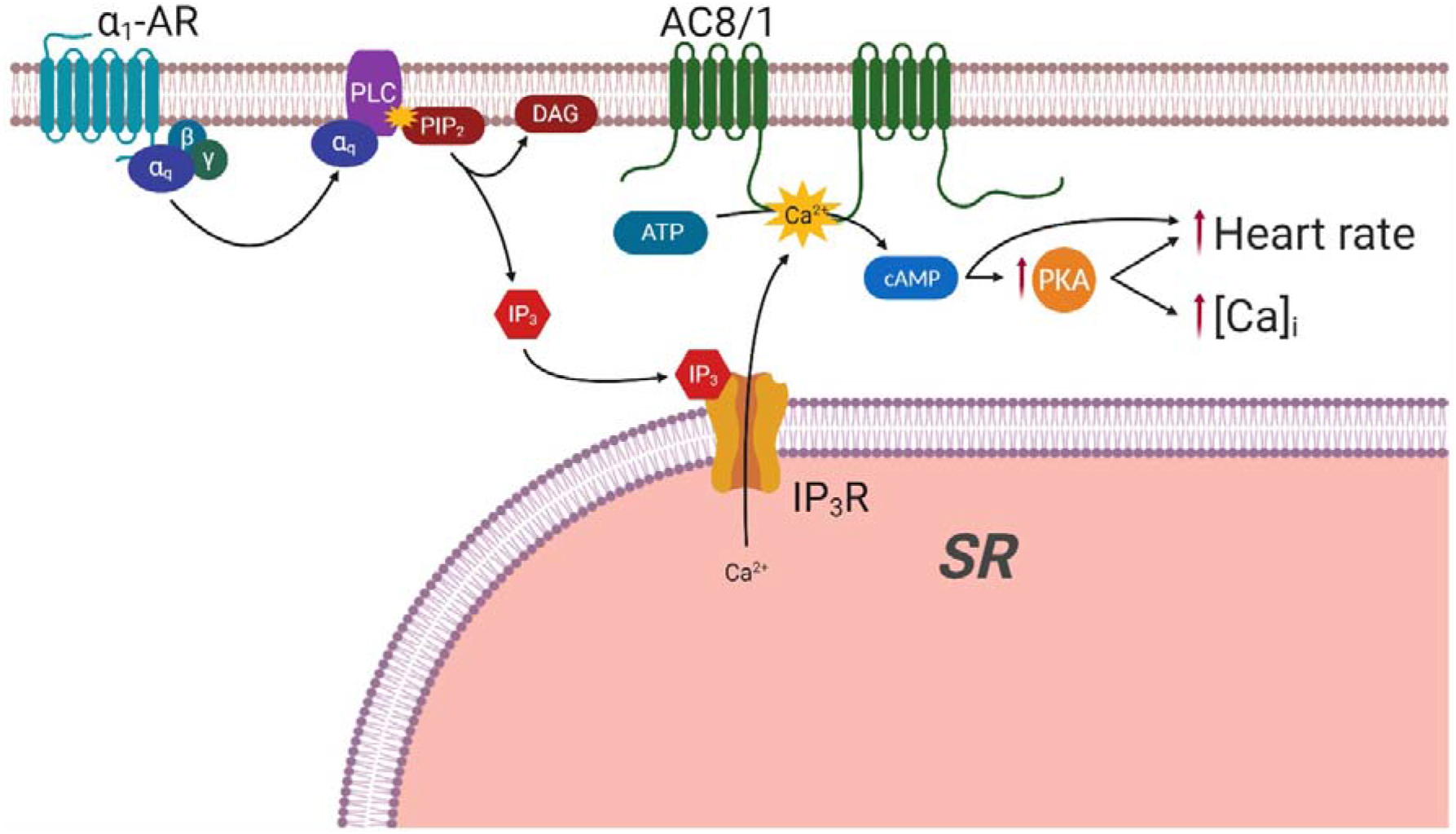
Proposed mechanisms for regulation of intracellular calcium via IP3 signalling. Scheme indicates potential mechanisms by which activation of α1-AR by PE may lead to increased atrial cytoplasmic calcium transients (indicated by [Ca^2+^]_i_) and sino-atrial node beating rate based on published observations in addition to our present results. Activation of α1-AR leads to elevated IP_3_ resulting from cleavage of PIP_2_ to DAG and IP_3_ by PLC. IP_3_ activation of IP_3_R_2_ results in release of Ca^2+^ from the SR, which subsequently leads to activation of Ca^2+^-sensitive adenylyl cyclase (AC8 or AC1) and activation of PKA by cAMP, or direct effects of cAMP on the funny current I(f). In the proposed scheme AC8 is placed in the sarcolemma, but it remains to be established whether there is an additional location in the junctional SR and whether nearby AC1 may also be activated by IP_3_-mediated Ca^2+^ release. Image created with BioRender.

Under the conditions of our study, inhibition of ACs or PKA significantly reduced baseline stimulated Ca^2+^ transient amplitude in atrial myocytes and beating rate in right atrial preparations. This is consistent with published data from our group ^17,18^ and others ^54,55^. Indeed, it has previously been shown that heart rate in AC8 over-expressing mice is significantly higher than in their wild-type counterparts ^56^. The sino-atrial node has a constitutive level of cAMP which is significantly greater than that of the ventricle in the absence of adrenergic stimulation ^57^. How much of this activity is attributable specifically to Ca^2+^-stimulated ACs is not discernable from our experiments as selective inhibitors are not currently available for all ACs. The diastolic cell Ca^2+^ concentration in SAN myocytes, ~225 nmol/L ^58^, is considerably higher than that in ventricular cells. Given that AC1 and AC8 proteins have not been shown to be expressed in ventricular tissue ^17^, it seems likely that cAMP production by Ca^2+^-stimulated ACs could contribute to the differences between resting SA nodal and ventricular cAMP concentration previously reported ^57^. Indeed, cAMP synthesis activity is high in SAN myocyte lysates in 1 μmol/L Ca^2+^ but almost abolished in Ca^2+^-free solution ^42^, suggesting Ca^2+^-stimulated cAMP production may be the dominant mechanism in these cells at rest. Our data indicate that high constitutive cAMP production in the atria and sino-atrial node cannot be attributed to background IP_3_R activity under the conditions of our experiments, as 2-APB alone did not have a significant effect on cellular Ca^2+^ transient amplitude or tissue beating rate.

In atrial myocytes isolated from cat, the effects of PE to enhance I_CaL_ have been reported to occur through inhibition of phosphodiesterase downstream of PI-3K-mediated eNOS activation ^24^. It was concluded that the primary role of IP_3_-mediated Ca^2+^ release in this process was to stimulate eNOS. Whilst we agree that cAMP and PKA are central to the response of atrial myocytes and the sino-atrial node to PE, and IP_3_, we did not find evidence that nitric oxide or soluble guanylyl cyclase activity was required for enhancement of atrial whole-cell Ca^2+^ transients in the guinea pig or SA-nodal beating rate in the mouse under the conditions of our experiments.

It has previously been hypothesised that Ca^2+^ release through IP_3_Rs acts to enhance atrial myocyte Ca^2+^ transients by increasing the local Ca^2+^ concentration around RyRs and therefore enhancing RyR response to the opening of LTCC ^1,14^. Although this is plausible, and has been observed in an IP_3_R over-expression model ^59^, our data investigating the direct effect of IP_3_ photorelease in the presence of downstream inhibitors, supported by that of Wang *et al.* (2005) ^24^ using indirect stimulation of the IP_3_ pathway, is consistent with the notion that the functional, physiological consequences of IP_3_R opening in cardiac atrial myocytes may result predominantly from modulation of other signalling pathways as opposed to direct effects on RyR responsiveness.

Even in healthy atrial myocytes, stimulation of IP_3_Rs can generate arrhythmogenic Ca^2+^ waves ^1,3^. In fact, stimulation of rat atrial myocytes demonstrated that IP_3_ is more arrhythmogenic than levels of either digoxin or endothelin producing a similar change in Ca^2+^ transient amplitude, and more arrhythmogenic than isoprenaline despite a greater Ca^2+^ transient response to this sympathomimetic ^3^. IP_3_R expression is significantly increased in cells from AF patients ^10^, and animal models of AF ^11^. Further, IP_3_R expression is increased in atrial myocytes from a heart failure model ^60^, a condition strongly associated with development of AF. In addition, Mougenot *et al.* (2019) ^61^ have recently demonstrated that overexpression of AC8 accelerates age-related cardiac dysfunction through increased hypertrophy and interstitial fibrosis in transgenic mice. This evidence highlights the importance of understanding the cellular mechanisms of the IP_3_ pathway and role of Ca^2+^-sensitive ACs in healthy and diseased cardiomyocytes.

The present paper provides novel information regarding the signalling pathways responsible for physiological responses to IP_3_, namely a crucial requirement for cAMP and PKA. In particular, we have focused on the position of Ca^2+^-stimulated ACs as an effector of this interaction. These novel data are not only interesting in that they provide an added level of complexity to Ca^2+^ modulation in the atria and sino-atrial node, but that they also raise questions about the mechanisms and role of this signalling in common pathology.

## Methods

### Atrial myocyte isolation

Male Dunkin Hartley guinea pigs (350-550g, Envigo or B&K Universal) were housed and maintained in a 12 h light-dark cycle with *ad libitum* access to standard diet and sterilised water. Guinea-pigs were culled by cervical dislocation in accordance with Home Office Guidance on the Animals (Scientific Procedures) Act (1986). Atrial myocytes were isolated following the method of Collins et al. (2011) ^62^ and stored at 4°C in a high potassium medium containing (in mmol/L): KCl 70, MgCl_2_ 5, K^+^ glutamine 5, taurine 20, EGTA 0.1, succinic acid 5, KH_2_PO_4_ 20, HEPES 5, glucose 10; pH to 7.2 with KOH. Healthy atrial myocytes were identified on the basis of morphology.

### Immunocytochemistry

Immunocytochemistry staining and analysis was carried out using the method of Collins and Terrar (2012)^18^. AC1 (sc25743) and AC8 (sc32128) primary antibodies were purchased commercially (Santa Cruz Biotechnology) and used at a dilution of 1:200. IP_3_R monoclonal primary antibodies (IP_3_R1 KM1112, IP_3_R2 KM1083, IP_3_R3 KM1082) were a kind gift from Prof Katsuhiko Mikoshiba ^63^ and used at a dilution of 1:1000. All primary antibody staining was carried out overnight at 4 °C. Secondary antibody labelling was carried out using either AlexaFluor −488 or −555 conjugated secondary antibodies (Invitrogen), raised against the appropriate species, for 60 minutes at room temperature at a dilution of 1:400. Observations were carried out using a Leica DMIRB inverted microscope modified for confocal laser-scanning microscopy (x63 water objective) and Leica TCSNT software or using a Zeiss LSM 510 (x40 oil objective). For detection of AlexaFluor 488, fluorescence excitation was at 488 nm with emission collected >515 nm. An excitation filter of 543 nm and an emission filter at 600 ± 15 nm were used to detect AlexaFluor 555. In order to quantify the relationship between the red and green signals that were created during double labelling experiments, we carried out a pixel-by-pixel co-localisation analysis. The analysis used produced Pearsons coefficient, which is between −1 (total exclusion of the signals) and +1 (complete co-localisation of the signals).

### Ca^2+^ transient imaging and IP_3_ Photorelease

For whole-cell fluorescence experiments, isolated atrial myocytes were incubated with Fluo-5F (3 μmol/L) for 10 min then plated to a glass cover slip for imaging. Carbon fiber electrodes were used to field-stimulate Ca^2+^ transients at a rate of 1 Hz. All experiments were carried out at 35 ± 2°C (fluctuation within a single experiment was <0.5°C) under gravity-fed superfusion of physiological salt solution (PSS, in mmol/L): NaCl 125, NaHCO_3_ 25, KCl 5.4, NaH_2_PO_4_ 1.2, MgCl_2_ 1, glucose 5.5, CaCl_2_ 1.8, oxygenated with 95 % O_2_ / 5 % CO_2_. Solution flow rate was 3 mL min^−1^.

For photorelease experiments, isolated atrial myocytes were incubated for 60 minutes at room temperature with 0.5 μmol/L membrane-permeant caged IP_3_ (caged-IP_3_/PM) and 0.025 % Pluronic F127 (Enzo Life Sciences). 3 μmol/L Fluo-5F-AM was added for the last 10 minutes of incubation. DMSO concentrations were 0.5 % during IP3/PM loading and 0.75 % during IP_3_/PM+Fluo-5F. Cells were visualised using a Zeiss Axiovert 200 with attached Nipkow spinning disc confocal unit (CSU-10, Yokogawa Electric Corporation). Excitation light, transmitted through the CSU-10, was provided by a 488 nm diode laser (Vortran Laser Technology Inc.). Emitted light was passed through the CSU-10 and collected by an iXON897 EM-CCD camera (Andor) at 65 frames per second. UV uncaging was carried out using 3x rapid flashes of a Xenon arc lamp (Rapp Optoelectronics), delivered through the objective lens. For inhibitor work, each aliquot of IP_3_/PM (3-4 experiments) was first used for a control experiment and inhibitor data were excluded if control cells did not respond. Cells were also excluded if, upon analysis, control (pre-photorelease) data exhibited alternans, missed beats or were otherwise unstable.

### Murine atrial studies

Adult male CD1 mice (30-35 g, Charles River UK CD-1® IGS) were housed maintained in a 12 h light-dark cycle with *ad libitum* access to standard diet and sterilised water. Mice were culled by cervical dislocation in accordance with Home Office Guidance on the Animals (Scientific Procedures) Act (1986). The heart was rapidly excised and washed in heparin-containing PSS. The ventricles were dissected away under a microscope and the area adjacent to the sino-atrial node cleared of connective tissue. The spontaneously beating atrial preparation was mounted in a 37 °C organ bath containing oxygenated PSS and connected to a force transducer (MLT0201 series, ADInstruments) in order to visualise contractions.

Resting tension was set between 0.2 and 0.3 g, the tension signal was low-pass filtered at 20 Hz and beating rate calculated from the time interval between contractions. After stabilisation (variation in average rate of a 10s sample of no more than 2 bpm over a 10-minute period), cumulative concentrations of PE were added to the bath (range 0.1 to 30 μmol/L) in the presence of metoprolol (1 μmol/L, applied 30 min prior to PE) to ensure specificity to α-adrenergic effects. Preparations were excluded if stabilised beating rate under control conditions (PSS only) was less than 300 bpm or if preparations were not rhythmic.

### Statistics

For all single cell data, t-tests or ANOVA were used as appropriate. Experimenters were not blinded to the conditions being analysed. Log(concentration)-response curves, used to estimate EC50s and maximum responses, were calculated using Prism8 software (GraphPad, CA, USA), by fitting an agonist-response curve with a fixed slope to normalised response data. Normalised data was used to compare responses as it was expected some inhibitors used would significantly affect the control beating rate or Ca^2+^ transient amplitude. Data are presented as mean ± SEM, other than EC50 which are presented as best-fit value with 95 % confidence interval.

## Additional Information

### Data Availability Statement

Please contact the corresponding author for all reasonable requests.

## Acknowledgements

RABB is funded by a Sir Henry Dale Wellcome Trust and Royal Society Fellowship (109371/Z/15/Z). This project was supported by a British Heart Foundation Project Grant (PG/18/4/33521). RAC is a post-doctoral scientist funded by the Wellcome Trust and Royal Society (109371/Z/15/Z). TPC was funded through a British Heart Foundation DPhil studentship (FS/05/121) in the DAT lab. SJB is a post-doctoral scientist funded by the British Heart Foundation (PG/18/4/33521). TA received funding from the Returners Carers Fund (PI RABB), Medical Science Division, University of Oxford, the Nuffield Benefaction for Medicine and the Wellcome Institutional Strategic Support Fund (ISSF), University of Oxford.

## Author Contributions Statement

RAC carried out IP3 photorelease work. TPC carried out immunofluorescence work. SJB produced Figure 4. RAC and SJB wrote the manuscript. RAC, TA and SJB carried out cell isolations. MZ contributed intellectually to study design and manuscript writing. RAC and SR carried out the atrial preparations. DAT, RAB and RAC designed the study. All authors have contributed to refinement of the manuscript.

